# Conformational landscape of the p28-bound human proteasome regulatory particle

**DOI:** 10.1101/088492

**Authors:** Ying Lu, Jiayi Wu, Shuobing Chen, Shuangwu Sun, Yong-Bei Ma, Qi Ouyang, Daniel Finley, Marc W. Kirschner, Youdong Mao

## Abstract

The proteasome holoenzyme is activated by its regulatory particle (RP) consisting of two subcomplexes, the lid and the base. A key event in base assembly is the formation of a heterohexameric ring of AAA-ATPases, which is guided by at least four RP assembly chaperones in mammals: PAAF1, p28/gankyrin, p27/PSMD9 and S5b. We determined a cryo-EM structure of the human RP in complex with its assembly chaperone p28 at 4.5-Å resolution. The Rpn1-p28-AAA subcomplex in the p28-bound RP is highly dynamic and was resolved to subnanometer resolution in seven states, which recapitulate the conformational landscape of the complex. Surprisingly, the p28-bound AAA ring does not form a channel in the free RP. Instead, it spontaneously samples multiple ‘open’ and ‘closed’ topologies. Our analysis suggests that p28 guides the proteolytic core particle to select certain conformation of the ATPase ring for RP engagement in the last step of the chaperone-mediated proteasome assembly.

## INTRODUCTION

The ubiquitin-dependent protein degradation, mediated by the proteasome, regulates numerous biological processes in all eukaryotes. The complete assembly of the 26S proteasome holoenzyme consists of a 20S proteolytic core particle (CP) and two 19S regulatory particles (RP) (Chen et al., 2016; Finley, 2009; Park et al., 2010; Tomko and Hochstrasser, 2013). Peptide cleavage is carried out inside the barrel-shaped CP chamber. The RP associates with the CP to activate its ubiquitin-dependent functions. A lid subcomplex was found to stably form prior to the completion of RP assembly in yeast (Glickman et al., 1998). The lid contains six subunits featuring Proteasome-Cyclosome-Initiation factor (PCI) domain (Rpn3, Rpn5-7, Rpn9, Rpn12), two subunits featuring MPR1/PAD1 amino-terminal (MPN) domain (Rpn8 and Rpn11) and a small tether protein Sem1 (Finley, 2009). The six PCI proteins are assembled into a horseshoe-like shape encircling the Rpn8-Rpn11 heterodimer (Beck et al., 2012; Chen et al., 2016; da Fonseca et al., 2012; Dambacher et al., 2016; Lander et al., 2012; Lasker et al., 2012; Luan et al., 2016; Sledz et al., 2013; Unverdorben et al., 2014). Rpn11, the only essential deubiquitylating enzyme in the proteasome, resides at the entrance to the substrate-translocation channel, and removes ubiquitin chains *en bloc* from a substrate during degradation(Pathare et al., 2014; Verma et al., 2002; Worden et al., 2014; Yao and Cohen, 2002). The rest of the RP subunits, apart from Rpn10, form a 9-subunit subcomplex named the base (Chen et al., 2016; Lander et al., 2012; Lasker et al., 2012). A critical component in the base is a peptide-unfolding channel made of hetero-hexameric AAA-ATPase ring, Rpt1-6. Each Rpt subunit features three domains, namely, coiled-coil (CC), oligonucleotide/oligosaccharide-binding (OB) and ATPase associated with diverse cellular activities (AAA) domains (Chen et al., 2016; Huang et al., 2016; Schweitzer et al., 2016; Zhang et al., 2009a; Zhang et al., 2009b). In the 26S holoenzyme, the AAA domains assemble into a helical staircase architecture and harvest the free energy from ATP hydrolysis to power substrate unfolding and translocation into CP chamber for degradation (Beck et al., 2012; da Fonseca et al., 2012; Lander et al., 2012).

A key event in base assembly is the formation of a heterohexameric ring of AAA-ATPases, which is guided by at least four RP assembly chaperones in mammals: PAAF1 (proteasomal ATPase-associated factor 1), p28/gankyrin (Krzywda et al., 2004; Nakamura et al., 2007a), p27/PSMD9 and S5b (Roelofs et al., 2009; Tomko and Hochstrasser, 2013). Each RP assembly chaperone binds a distinct base subunit, forming three RP assembly precursors, Rpt3-Rpt6-p28-PAAF1, Rpt4-Rpt5-p27, and Rpt1-Rpt2-S5b complexes (Kaneko et al., 2009; Park et al., 2009; Roelofs et al., 2009). Release of chaperones p28, PAAF1, and S5b upon holoenzyme assembly is important for proteasome activation. The yeast ortholog of p28 is Nas6. Modeling of the Rpt3-Nas6 structure (Nakamura et al., 2007a; Nakamura et al., 2007b), into an ATPase ring of the proteasome holoenzyme suggests that Nas6 physically occludes the formation of proper RP-CP contacts (Park et al., 2013; Roelofs et al., 2009). To date, no structural information is available for a complete RP in complex with any assembly chaperones. It remains unclear how RP chaperones interact with the RP to ensure proper complex assembly, and how the chaperones are released to activate the proteasome. To address these problems, we attempted to analyze the structures of the free RP in complex with its chaperone p28 by single-particle cryo-electron microscopy (cryo-EM).

## RESULTS

### Structural Analysis

We purified the p28-bound RP from human embryonic kidney (HEK) 293 cells, using biotin-tagged Rpn11, followed by size-exclusion chromatography (Figure 1A, B, Figure S1) (Guerrero et al., 2006). Through exhaustive computational classification of single-particle cryo-EM data (Figures S2 and S3), we identified a homogeneous dataset of the free RP, which was refined to a nominal resolution of 4.5 Å by a gold-standard procedure, in which two half-datasets were refined separately (Figure 1C). The high-resolution components include all the RP components but Rpn1, Rpn13, and the AAA domains of the ATPases. An atomic model was built and refined based on this density map, which includes the complete lid, Rpn2, Rpn10, and the CC (coiled coil) and OB (oligonucleotide/oligosaccharide-binding) domains of Rpt1-Rpt6 (Figure 1D, Figure S4 and Table S1).

**Figure 1.**
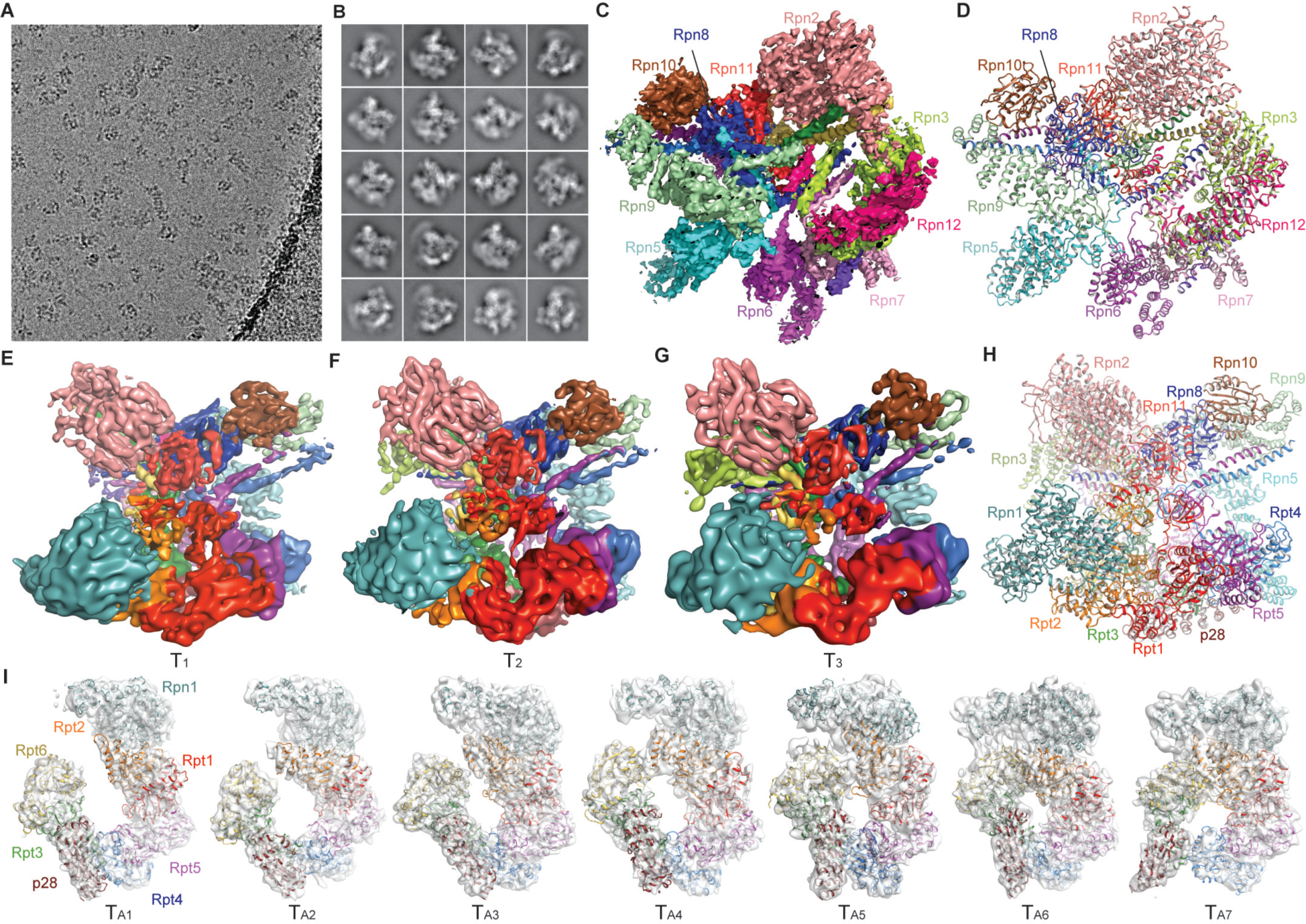
Cryo-EM analysis of the human RP subassemblies. (A) Typical cryo-EM micrograph showing the projections of the free human RP. (B) A gallery of representative reference-free class averages of the free human RP. (C) High-resolution cryo-EM density map of the free RP that allows the atomic modeling. (D) The atomic model refined against the cryo-EM density shown in panel (C). (E)-(G) The cryo-EM density maps of the conformational states T_1_ (E), T_2_ (F) and T_3_ (G). (H) Hybrid atomic model of the free RP, where the atomic model of the non-AAA component is combined with the pseudo-atomic model of AAA domains of Rpt1-6. (I) Pseudo-atomic models of Rpn1-p28-AAA components in cartoon representation superimposed with the corresponding cryo-EM density in transparent surface representation. The map-superposed models correspond to the seven states T_A1-7_ and are aligned into a sequence that following the decrease of the Rpt2-Rpt6 gap from T_A1_ to T_A7_, and from the left to the right.

Through 3D classification, we obtained three conformational states of the complete p28-bound free RP, designated T_1_, T_2_ and T_3_, differing mainly in their AAA domains of the ATPase at 5.2 Å, 6.1 and 6.8 resolution, respectively (Figure 1E-G and Figure S5A-C). The AAA and CC-OB domains of each Rpt subunit are connected through a flexible loop that allows inter-domain motions. In the cryo-EM image data, the movement of the entire hexameric AAA ring relative to the OB ring is coupled with the inter-domain motion between adjacent Rpt AAA domains in the free RP. As a result, Rpn1, p28, and the AAA domains of Rpt3-Rpt6 exhibit limited resolution and diffuse densities in the cryo-EM maps of the complete free RP. To decouple the two dynamic modes, we employed a ‘density-subtraction’ strategy in which the density of the non-AAA subcomplex is subtracted from each single-particle image. Maximum-likelihood-based classification (Scheres, 2012a, b) of the density-subtracted images allowed us to obtain seven conformational states of the Rpn1-p28-AAA subcomplex, designated T_A1-7_, each refined separately to 7-9 Å resolution (Figure 1I, Figures S3 and S5D-J). The cryo-EM maps of these Rpn1-p28-AAA conformations significantly improved the local resolution in Rpn1, p28/gankyrin, and the AAA domains of Rpt3-Rpt6. Based on the atomic structure of the human proteasome holoenzyme (Chen et al., 2016), we built pseudo-atomic models for the Rpn1-p28-AAA subcomplex in the T_A1-7_ states. Integrating these pseudo-atomic models with the atomic model of the non-AAA components allowed us to generate hybrid atomic models for the complete p28-RP complexes in the T_A1-7_ states (Figure 1H).

### Structures of the p28-Bound RP

The cryo-EM densities of the non-ATPase components in the T_1_, T_2_ and T_3_ states show virtually the same conformation as that in the 4.5-Å map of the non-AAA subcomplex (Figure 2A-C). The T_1_, T_2_ and T_3_ structures contain six Rpt subunits. The local resolutions are lower in the AAA domains than in the OB domains. The ATPase heterohexamers in both T_1_ and T_2_ states exhibit a prominent opening between Rpt2 and Rpt6 (Figure 2D), whereas Rpt2 and Rpt6 are in contact in the T_3_ state (Figure 2E, F). The N-terminal elements of Rpn2, Rpn3 and Rpn12 that form a trimeric interface at the apex of RP also suffer diffuse cryo-EM densities in the T_1_ state indicative of local conformational dynamics (Figure 2A), whereas corresponding local densities in the T_2_ and T_3_ states are substantially stronger (Figure 2B,C).

**Figure 2.**
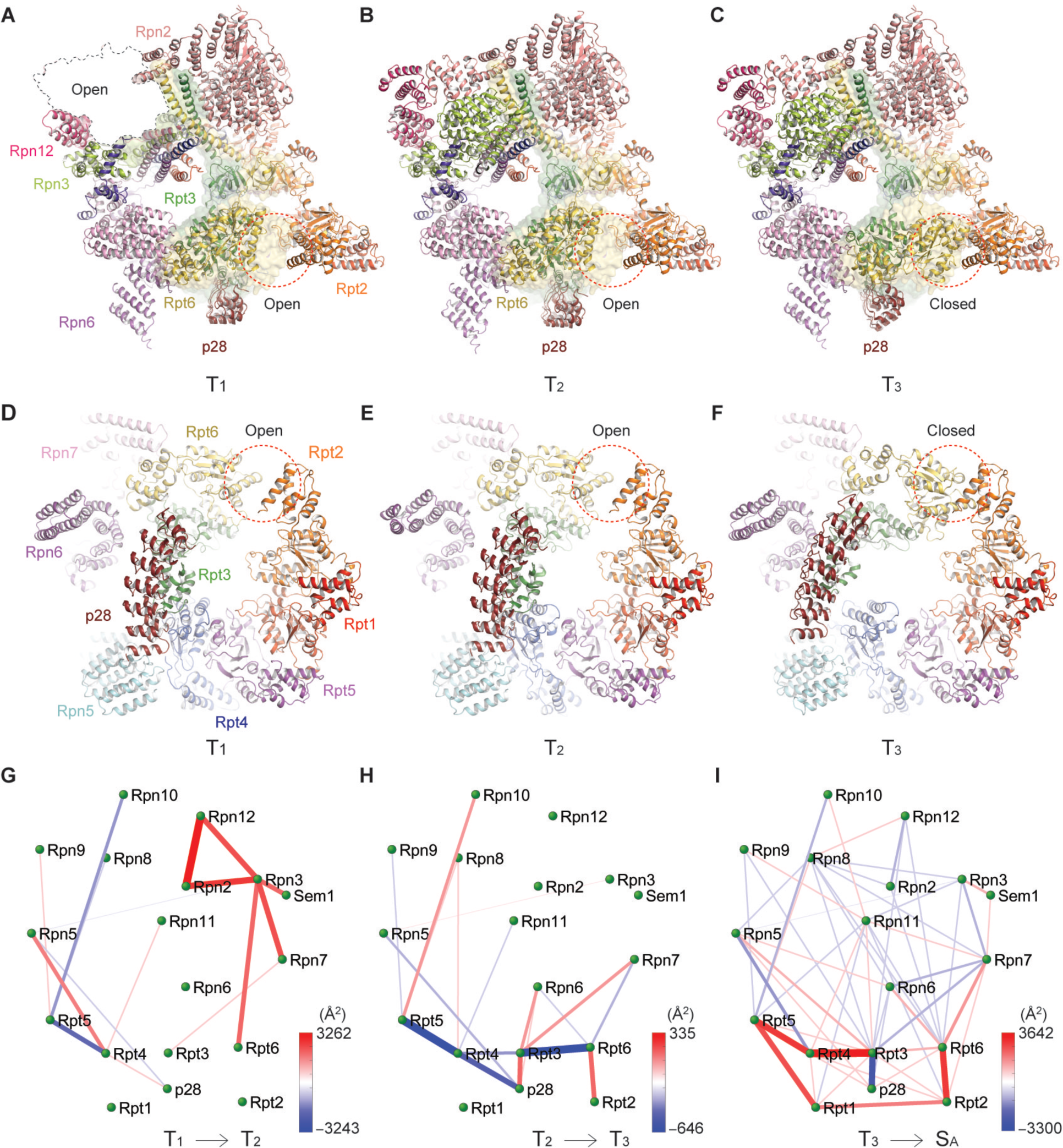
Transitions between RP assemblies in different conformations. (A)-(C) The pseudo-atomic model of the free RP in the T_1_ (A), T_2_ (B) and T_3_ (C) states. (D)-(F) Side-by-side comparison of the three RP structures in the T_1_ (D), T_2_ (E) and T_3_ (F) states from the perspective rotated by 90° relative to (A) to (C). (G)-(I) The microinteractome networks visualize the changes of inter-subunit interface areas during the four transitions, from T_1_ to T_2_ (G), from T_2_ to T_3_ (H), from T_3_ to S_A_ (I).

To capture the ‘hotspots’ of inter-subunit interface changes upon structural transitions, we calculated the differences of inter-subunit interfaces and mapped them into differential networks of inter-subunit interactions, termed ‘microinteractome networks’ (Figure 2G-I). The T_1_-to-T_2_ transition features a ‘hotspot’ around Rpn3 (Figure 2G). In the T_2_-to-T_3_ transition, an increased Rpt2-Rpt6 interaction trades off for a decreased interaction between gankyrin and Rpt4 (Figure 2H). The closure of Rpt2-Rpt6 gap is accompanied with increased Rpt3 interactions with Rpn6 and Rpn7. The T_3_-to-S_A_ transition network suggests subtle inter-subunit rearrangement in the non-ATPase components, in contrast to dramatic strengthening of inter-Rpt interactions upon RP-CP association (Figure 2I).

### Conformational Changes of the Non-AAA Subcomplex upon RP-CP Association

Comparison of the free RP structure with that of proteasome holoenzyme in a substrate-accepting (S_A_) state (Chen et al., 2016) reveals modest conformational changes of the lid upon RP-CP association (Figure 3A-D). When the free RP structure is aligned with the holoenzyme structure based on their lid subcomplexes (Figure 3A), the N-terminal PCI domain of Rpn7 is translated ~15 Å towards the CP, due to rearrangement of Rpt3 and Rpt6 during RP-CP association. The most prominent conformational change is observed in Rpn6. The N-terminal PCI domain of Rpn6 exhibits a dramatic rotation of 40°, allowing Rpn6 to interact with α2 subunit of the CP (Figure 3A). When the subunit structures are aligned separately on their own, there is a ~40° rotation of the C-terminal helix around a pivot point at residue Pro317 in Rpn6 (Figure 3B), whereas a ~20° rotation of the C-terminal helix is seen in Rpn7 around a pivot point at residue Pro359 (Figure 3C). The rest of the lid subunits, as well as Rpn2 and Rpn10, are nearly identical in both structures (Figure 3D).

**Figure 3.**
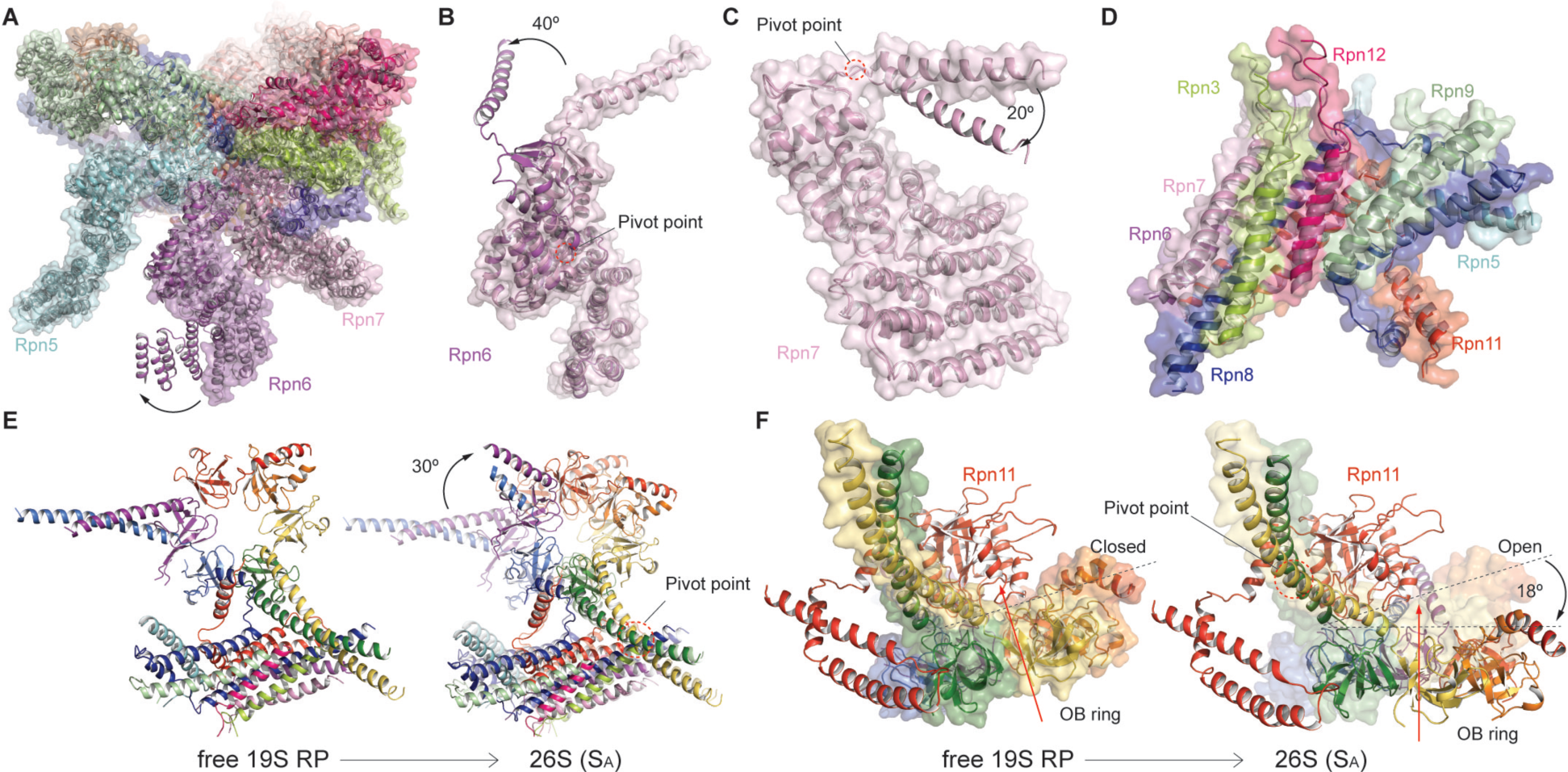
Conformational transitions of the non-AAA subcomplex upon 26S assembly. (A) Superposition of the structures of the free RP with the RP in 26S in the S_A_ state, after aligning them over the non-AAA subcomplex. (B)-(D) Superposition of the structures of Rpn6 (B), Rpn7 (C), and the lid central helical bundle (D), by alignment their own structure in the free RP with that in the 26S. (E) Side-by-side comparison of the RP components including CC-OB and lid helical bundle, showing a 30° in-plane rotation between the free RP in the T_3_ state and the 26S in the S_A_ sate. On the right, the T_3_ structure is rendered in transparent cartoon for convenience of comparison. (F) Side-by-side comparison of the RP components including CC-OB and lid helical bundle, showing a 18° out-of-plane rotation between the free RP in the T_3_ state and the 26S in the S_A_ sate. On the right, the T_3_ structure is rendered in transparent surface for convenience of comparison.

Although the structure of the AAA ring in the free RP is significantly different from that in the assembled proteasome, the OB ring in the free RP is nearly identical to that in the holoenzyme (Chen et al., 2016). Upon RP-CP assembly, the OB ring rotates in-plane by 30° and tilts out-of-plane by 18° (Figure 3E, F). The gap between Rpn11 and the OB ring is slightly narrower in the free RP than in the SB, SC and SD states of the proteasome holoenzyme (Chen et al., 2016), suggesting that Rpn11 tightly blocks the substrate entry port in the free RP (Figure 3F, Figure S6A-D). Indeed, an unfolding assay used in this study to characterize the free RP suggests that it essentially lacks substrate unfolding activity toward a fluorescent model substrate (see below and Figure 6). The free RP exhibits certain ATP-independent deubiquitiylating activity that is lower than that of the proteasome holoenzyme (Figure S7). In line with this observation, docking of the ubiquitin structure to the surface of Rpn11 in the free RP indicates that the active site for deubiquitylation is exposed on the molecular surface (Figure S6E, F). Conformation of the Rpn11-OB interface in a SB, SC or SD-like state has been suggested to enhance Rpn11 activity (Chen et al., 2016; Pathare et al., 2014; Worden et al., 2014), which may also account for the deubiquitylating activity of the free RP.

### Microstates of Rpn1-p28-AAA Subcomplex

A proper hexamerization of the AAA domains of Rpt1-6 is crucial for activating the substrate unfolding activity of the RP in an assembled proteasome holoenzyme. The cryo-EM structures of the Rpn1-p28-AAA subcomplex in seven states T_A1-7_ exhibit dramatic motions in their p28-Rpt3-Rpt6 and Rpn1 components (Figure 4A, Figure S5). The AAA domains form a C-like shape with an opening between Rpt2 and Rpt6 in all but the T_A6_ and T_A7_ states. The width of the opening is different in each state, and we used this feature to order the states from the widest to the tightest (Figure 4A), revealing a sequence of motion of the AAA ring through consecutive intermediates in an open-to-closed transition. Consistent with our assignment of the state sequence, the microinteractome networks of inter-subunit interactions exhibit a gradually increased connectivity from T_A1_ to T_A7_ (Figure 4B). Remarkably, Rpn1 rotates as the Rpt2-Rpt6 gap narrows. Rpn1-Rpt2 interaction is established starting from the T_A5_ state, suggesting that Rpn1 assists the open-to-closed transition. Correspondingly, Rpt3-bound p28 rotates in-plane by ~60° and out-of-plane by ~30° (Figure 4C). Accompanying closure of the Rpt2-Rpt6 gap, the Rpt3-Rpt4 interface is opened in the T_A7_ state, whereas in other states p28 appears to bridge the opening between Rpt3 and Rpt4.

**Figure 4.**
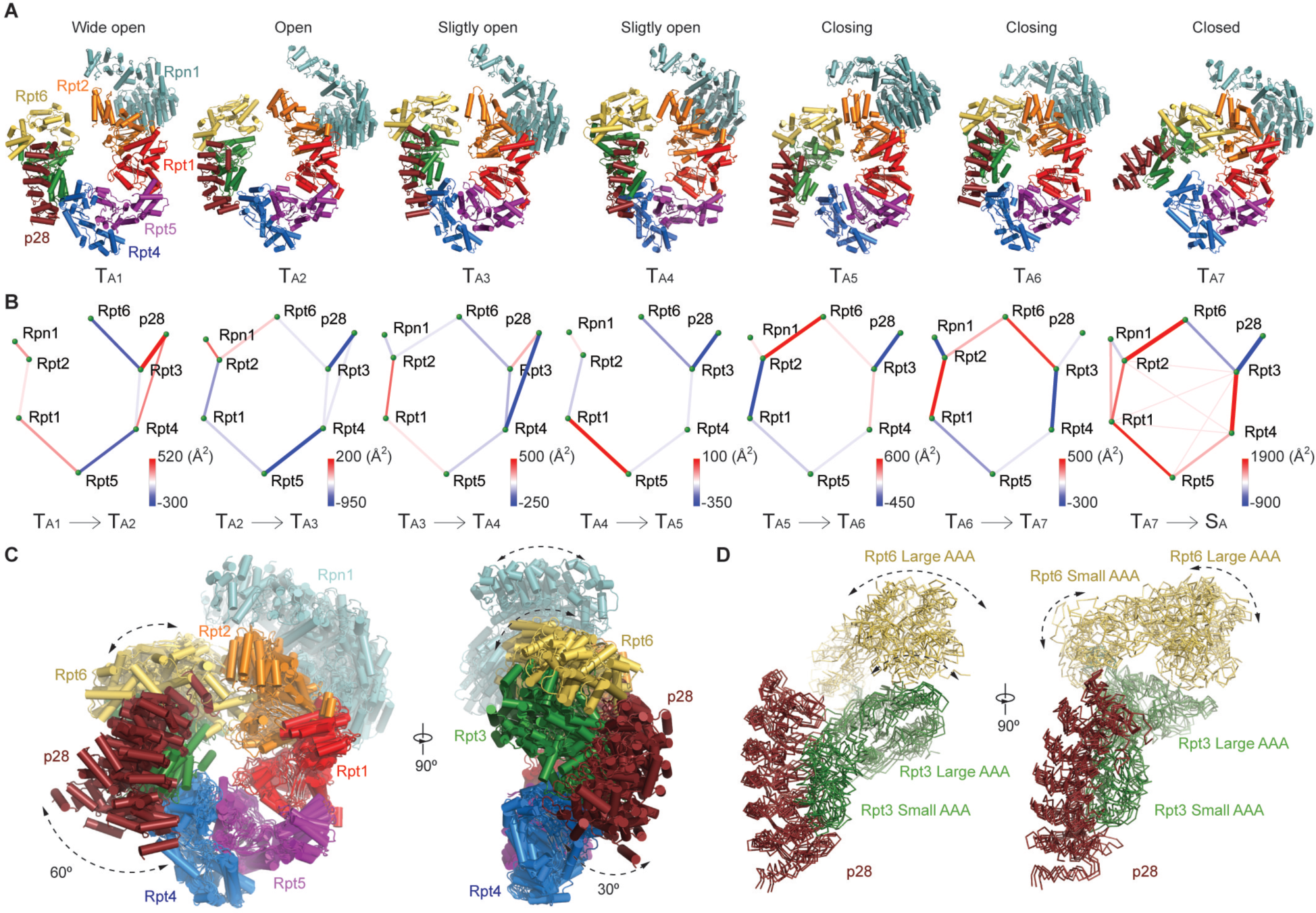
Conformational states of the Rpn1-p28-AAA components in the free RP. (A) The seven pseudo-atomic models of Rpn1-p28-AAA subcomplex from a viewing perspective that shows a large in-plane rotation of the p28/gankyrin. (B) The seven differential networks show the changes of inter-subunit interactions during the state transitions. (C) The seven pseudo-atomic models of the Rpn1-p28-AAA subcomplex are superimposed. On the right, a viewing perspective orthogonal to the left is shown. (D) The seven states are aligned relative to the p28-Rpt3 structure and superimposed with ribbon representations.

The elements that interact with p28 seem to undergo multiple modes of conformational dynamics (Figure 4C, D). The p28 movement orthogonal to the hexameric axis of the ATPase concurs with its wobbling along the axis (Figure 4C). Similar multi-modal remodeling at the Rpt2-Rpt6 interface is also observed (Figure 4D). Aligning the structures of p28-Rpt3 in the seven states reveals that the p28-Rpt3 interface is much more rigid than the Rpt3-Rpt6 interface (Figure 4D). Each AAA domain contains a large N-terminal α/β subdomain and a small C-terminal α-helical subdomain, separated by a short flexible linker. The small and large AAA subdomains exhibit different amplitudes of rotational motion (Figure 4D). In summary, the multi-modal dynamics of the AAA domains is characteristic of the p28-bound RP intermediate assembly.

### RP Remodeling upon RP-CP Association

To understand how the p28-RP approaches the CP for proteasome activation, we docked the pseudo-atomic model of the p28-RP complex in the T_A7_ state onto the CP, by aligning the AAA-ATPase in the p28-RP structure with that in the holoenzyme structure in the S_A_ state. Notably, the p28 protein shows minimal clashes with the α-ring of CP, Rpn6 or Rpn5 (Figure 5A), suggesting that p28 does not necessarily block the RP-CP association in its first-encounter state when the CP engages with a p28-bound RP in the T_A7_ state (Figure 5A-C). By contrast, there are prominent clashes between α2-subunit and p28 in the other six states, T_A1-6_ (Figure S8A). This suggests that p28 helps the CP to select the T_A7_ conformation to facilitate the RP-CP association by increasing interface contact area while introducing no significant occlusion (Figure 5D). Comparison of the docked p28-RP-CP intermediate complex with the holoenzyme structure reveals that the lid subcomplex undergoes a 40° rotation along an axis that is tilted ~30° from the heptameric axis of the CP (Figure 5E).

**Figure 5.**
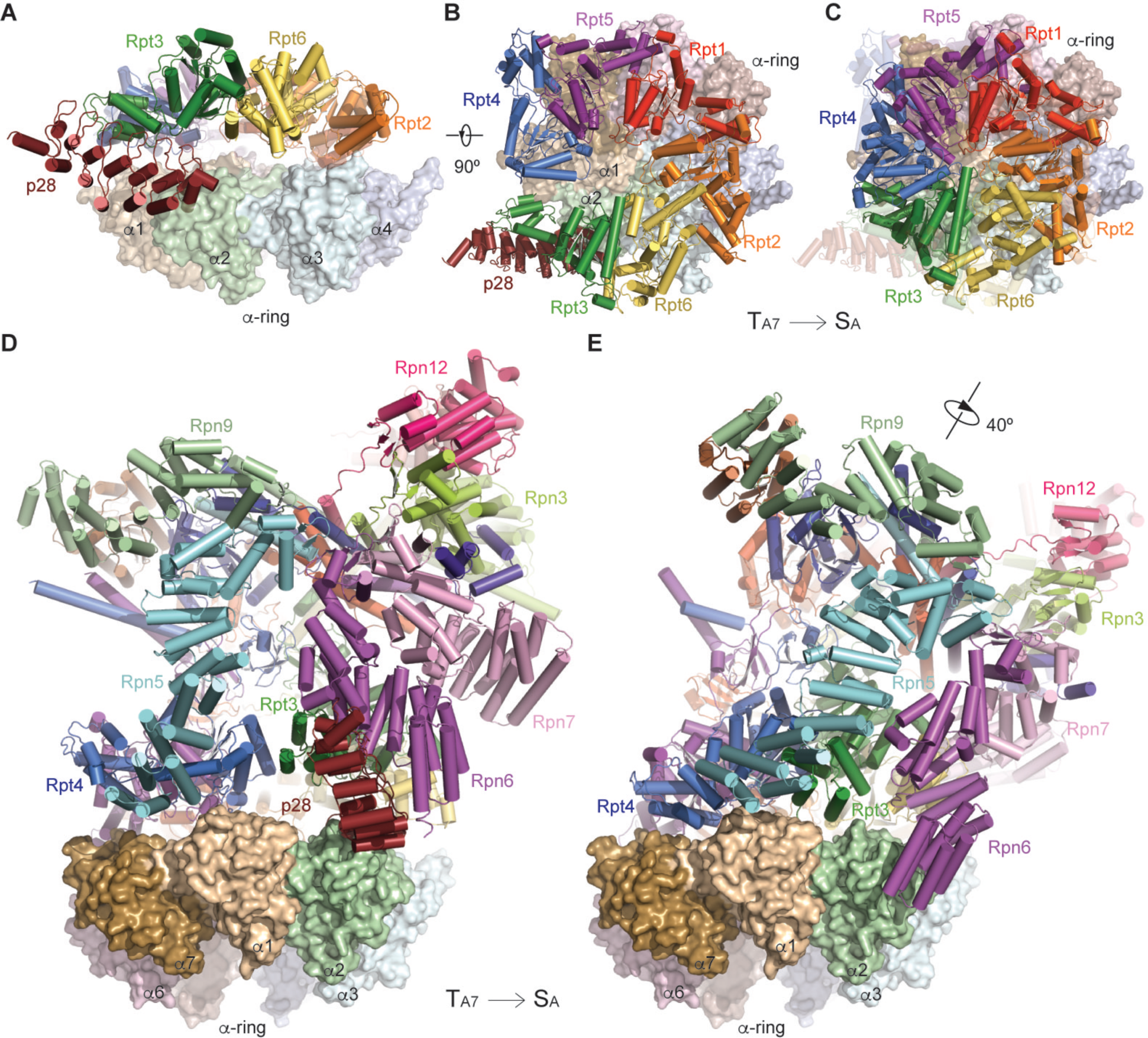
RP Remodeling upon RP-CP association. (A) The side view of the p28-AAA subcomplex in the T_A7_ state docking onto the α-ring surface, by aligning the AAA domains in the free RP against those in the 26S in the S_A_ state. The model predicts that p28 is subject to little or no clashes with the α-ring surface when the free RP first encounters the CP. (B) The top view of the p28-AAA subcomplex docking onto the α-ring surface. (C) The top view of the AAA hexamer on the α-ring surface in the 26S structure for comparison. The structure of p28-AAA subcomplex is superposed as a transparent cartoon. (D)The side view of the hybrid atomic model of the free RP in the T_A7_ state docking onto the α-ring surface, with the same alignment as used in panel (A). (E) The side view of the 26S in the S_A_ state, in the same orientation of the CP, showing that a large RP rotation accompanying the release of p28.

The assembly chaperone p28 has to be released upon the RPCP assembly and activation of proteasome (Roelofs et al., 2009). Our structural analysis suggests a shoehorn-like mechanism for p28 release. Although p28 can be an integral part of the RP-CP first-encounter complex, RP-CP association has to be reinforced with considerable remodeling of the AAA-ATPase and lid subcomplexes as well as the RP-CP interface. The stronger affinity of the α-ring with the ATPases, the closure of the gap between Rpt3 and Rpt4, and a tighter arrangement of the ATPase hexamer, may all help to drive p28 dissociation from Rpt3. Importantly, Rpt3 is translated ~15 Å upon completion of proteasome assembly to close its gap with Rpt4, which dramatically reorients its p28 binding site toward the α1-α2 surface (Figure 5B, C). Consistent with this picture, a mutant *rpt3-Δ1* was observed to be unable to release p28, perhaps because the mutated Rpt3 loses the affinity of its C-terminal tail with α-ring and is unable to complete the Rpt3 remodeling that ejects the assembly chaperone (Park et al., 2009).

### The Free RP is Deficient in Substrate Unfolding

To test whether a substrate translocation occurs during Rpn11-dependent deubiquitylation in the free RP, we developed a fluorescent-based, ubiquitin-dependent substrate-unfolding assay. As noticed previously, proteins fused with a green florescent protein (GFP) or its derivatives tend to be poor substrates of purified proteasomes, likely due to inefficient substrate unfolding and lack of proteasome turnover (Matyskiela et al., 2013). We found that a Sic1-fused, phytochrome-based near-infrared fluorescent protein (iRFP) was degraded efficiently upon ubiquitylation (Figure S7) (Shcherbakova and Verkhusha, 2013). We used it as a fluorescent reporter to investigate whether translocation occurs during the RP-driven deubiquitylation, since a loss of fluorescent signal will ensue due to threading through the ATPase channel as protein unfolding. To prevent refolding of iRFP, we employed a GroEL mutant that traps unfolded peptides and has been used in the study of GFP unfolding by ClpA (Weber-Ban et al., 1999). In contrast to the fluorescence loss due to proteasome-induced unfolding and translocation of the ssrA-fused iRFP, we did not observe any fluorescence signal change of ubiquitylated Sic1-iRFP, in the presence of the free RP and excess GroEL, suggesting lack of substrate unfolding activity (Figure 6). This is consistent with the observation that the Rpt AAA domains in the free RP form an open ring topology and are not yet assembled into a functional channel capable of substrate translocation.

**Figure 6.**
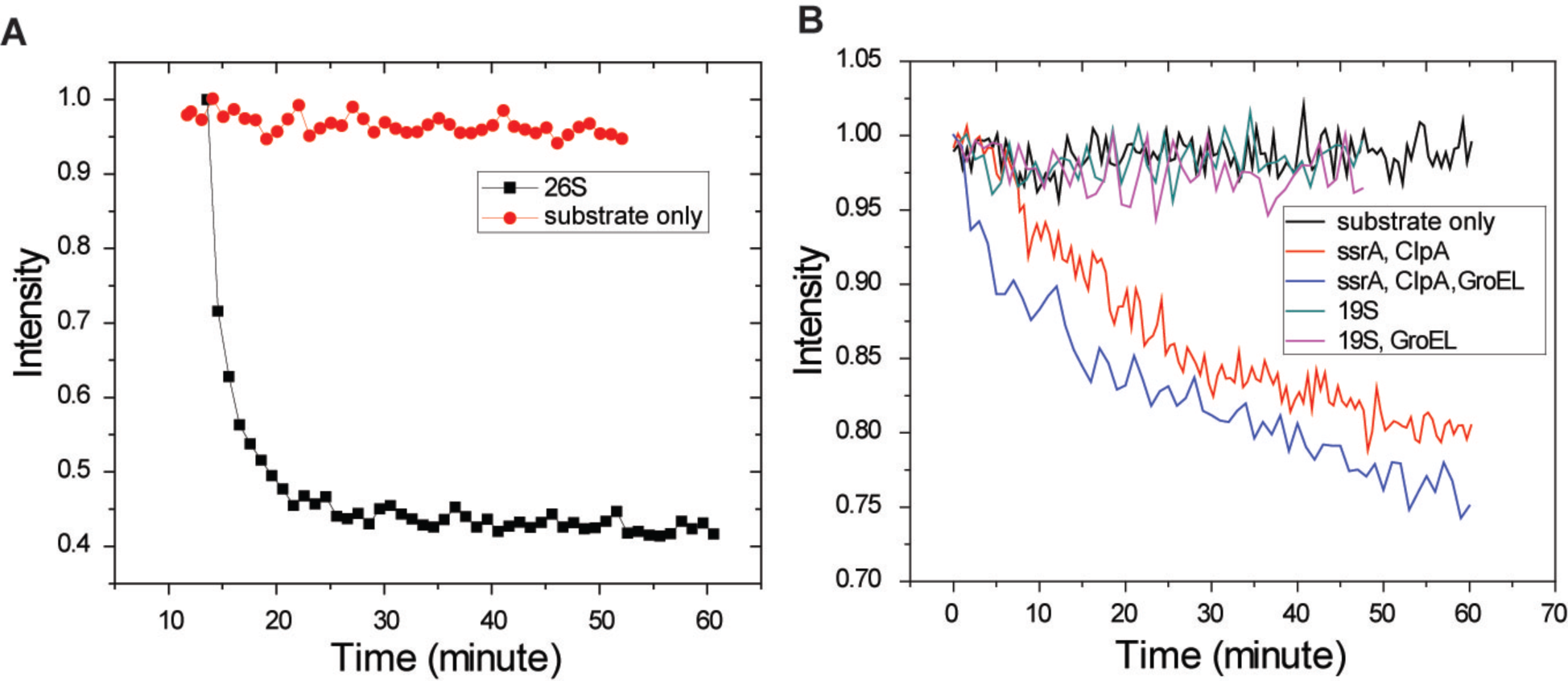
Fluorescent unfolding assay to examine free RP activity. (A) Ubiquitylated iRFP as a fluorescent reporter for proteasomal activity. 500nM Sic1-iRFP was ubiquitylated, purified and mixed with the 26S proteasome from yeast as described in the methods. Fluorescent intensity of the reporter was continuously monitored in a fluorescence spectrophotometer at room temperature. (B) Free RP lacks ATP-dependent unfolding activity. 500nM ubiquitylated Sic1-iRFP was mixed with the RP (19S) from yeast, in the presence or absence of the GroEL trap as described in methods. As a positive control, Sic1-iRFP carrying a C-terminal ssrA sequence (ssrA) was incubated with purified ClpA.

## DISCUSSION

To ensure a correct assembly of the heterohexameric AAA-ATPase, several assembly chaperones are involved in the RP assembly process to regulate the assembly sequence or rectify mis-assembled complexes (Kaneko et al., 2009; Park et al., 2009; Roelofs et al., 2009). The structures of the T_1-3_ subassemblies characterize an overall conformational landscape along the proteasome assembly pathway, hypothetically representing states of the ATPase hexamerization during its last step prior to the completion of holoenzyme assembly. In the absence of interactions with CP, the ATPase ring is unable to assume a stable rigid structure but instead samples a variety of conformations. Its conformational landscape was captured by seven microstates T_A1-7_ in the Rpn1-p28-AAA subcomplex. The approximately equal population of the seven states of the Rpn1-p28-AAA subcomplex suggests that each state may be transiently sampled by the free RP along the trajectory traversing the frustrated energy landscape of the complex (Wensley et al., 2010). This is consistent with the fact that they assemble into consecutive snapshots recapitulating an open-to-closed transition of the AAA ring. Our analysis suggests that the T_A7_ state of the AAA ring may facilitate RP-CP association and likely subsequent p28 release in a shoehorn-like fashion. Thus, p28 reconciles the remarkable structural differences in the RP before and after proteasome assembly, with guiding the CP to choose specific conformation of the ATPase ring for RP engagement, which completes the last step in the chaperone-mediated proteasome assembly.

A prominent feature of the RP assemblies is the blocking of substrate-entry port by Rpn11, which renders the OB-ring incompetent in accepting substrates. Moreover, the AAA channel is not yet formed in the free RP and undergoes strong conformational dynamics that spontaneously sample both open and closed ring topologies (Figure 4). The ATP-independent deubiquitylating activity of the free RP is detected in the absence of complete substrate unfolding in a fluorescent unfolding assay (Figure 6). Unfolded proteins, in the absence of degradation, could form aggregates impairing numerous cellular functions. The ‘closed’ OB-ring and ‘open’ AAA-ring in the free RP minimizes this risk prior to its incorporation into the 26S proteasome holoenzyme.

The overall conformation of the non-AAA components is virtually identical to that in the SB, SC and SD states of the human proteasome holoenzyme (Figure S6A, B) and resembles the yeast counterpart in the s2, s3 and substrate-engaged states (Chen et al., 2016; Luan et al., 2016; Matyskiela et al., 2013; Sledz et al., 2013; Unverdorben et al., 2014). This suggests that the non-AAA structure in the free RP could represent a ground state of this subcomplex. Indeed, the total buried interfacial area of the non-AAA subcomplex is about 900-Å^2^ larger in the free RP than in the proteasome holoenzyme in the S_A_ state (Chen et al., 2016). The non-AAA conformation of the proteasome in the S_A_ state is of higher potential energy and is stabilized by an additional interaction between Rpn5/Rpn6 and the α ring of the CP, which buries ~600-Å^2^ of interfacial area (Chen et al., 2016; Huang et al., 2016; Schweitzer et al., 2016). Taken together, our data suggest that the non-AAA subcomplex conformation switches mainly between its ground state and its higher-energy state during and after proteasome assembly (Chen et al., 2016; Luan et al., 2016; Matyskiela et al., 2013; Sledz et al., 2013; Unverdorben et al., 2014). The energetic proximity of the two states in the non-AAA subcomplex poises the RP adjacent to a ‘critical’ point that facilitates allosteric regulation of proteasome activation and function (Chen et al., 2016).

## EXPERIMENTAL PROCEDURES

### Protein Expression and Purification

Human 19S RP was affinity-purified on a large scale from a stable HEK293 cell line harboring HTBH tagged hRPN11 (a gift from L. Huang) (Wang et al., 2007). The cells were Dounce-homogenized in lysis buffer (50mM NaH_2_PO_4_ [pH7.5], 100mM NaCl, 10% glycerol, 5mM MgCl_2_, 0.5% NP-40, 5mM ATP and 1mM DTT) containing protease inhibitors. Lysates were cleared, then incubated with NeutrAvidin agarose resin (Thermo Scientific) overnight at 4°C. The beads were then washed with excess lysis buffer followed by the wash buffer (50mMTris-HCl [pH7.5], 1mM MgCl_2_ and 1mM ATP). Usp14 was removed from proteasomes using wash buffer +150mM NaCl for 30 min. Human 19S RP was eluted from the beads by cleavage, using TEV protease (Invitrogen), and was further purified by running gel filtration on a Superose 6 10/300 GL column at a flow-rate of 0.15ml/minute in buffer (30mM Hepes pH7.5, 60mM NaCl, 1mM MgCl_2_, 10% Glycerol, 0.5mM DTT, 0.8mM ATP). Gel-filtration fractions were concentrated to about 2 mg/ml. Right before cyro-EM sample preparation, the 19S RP sample was buffer-exchanged into 50mM Tris-HCl [pH7.5], 1mM MgCl_2_, 3mM ATP, 0.5mM TCEP, to remove glycerol, and was supplemented with 0.005% NP-40.

### Deubiquitylation and Degradation Assay

About 500nM ubiquitylated pySic1-iRFP was mixed with purified 19S RP or 26S proteasome in a buffer (50mM Tris-HCL [pH 7.5], 50mM NaCl, 5mM MgCl_2_, 0.5mM DTT, 0.2mg/ml BSA) supplemented with 3mM ATP or other nucleotides, incubated at room temperature. pySic1-iRFP was affinity purified using a Ni-NTA column after ubiquitylation.

### Substrate-Unfolding Assay

500nM purified, ubiquitylated pySic1-iRFP substrate was mixed with 20nM 19S RP or recombinant ClpA in a buffer (50mM Hepes [pH7.5], 20mM MgCl_2_, 10mM ATP, 150mM NaCl, 10% glycerol, 0.5mM DTT), in the presence or absence of GroEL trap at 500nM, continuously monitored in a Cary Eclipse fluorescence spectrophotometer (ex680/em715) at room temperature. In the fluorescent degradation assay, 20nM human 26S proteasome is employed instead of RP. The signal is normalized by the initial fluorescence intensity.

### Data Collection

A 2.5-μl drop of 3 mg/ml 19S solution was applied to a glow-discharged C-flat grid (R 1/1, 400 Mesh, Protochips, CA, USA), blotted for 2 second at 4 °C and 100% humidity, then plunged into liquid ethane and flash frozen using the FEI Vitrobot Mark IV. The cryo-grid was imaged in an FEI Tecnai Arctica microscope, equipped with an Autoloader, at a nominal magnification of 54,000 times and an acceleration voltage of 200 kV. Coma-free alignment was manually conducted prior to data collection. Cryo-EM data were collected semi-automatically by the Leginon version 3.1 (Suloway et al., 2005) on the Gatan K2 Summit direct detector camera (Gatan Inc., CA, USA) in a counting mode, with 9.0 s of total exposure time and 250 ms per frame. This resulted in movies of 36 frames per exposure and an accumulated dose of 50 electrons/Å^2^. The calibrated physical pixel size is 0.98 Å, respectively. The defocus in data collection was set in the range of from −1.0 to −3.0 μm. A total of 20,000 movies were collected, among which 16,111 movies were selected for further data analysis.

### Cryo-EM Data Processing and Reconstruction

The raw 36 movie frames were first corrected for their gain reference and each movie was used to generate a micrograph that was corrected for sample movement and drift. These drift-corrected micrographs were used for the determination of actual defocus of each micrograph with the CTFFind3 program (Mindell and Grigorieff, 2003). Particles were initially automated picked in SPIDER (Shaikh et al., 2008). 250,251 particles were manually checked from 16,111 micrographs in EMAN2 (Tang et al., 2007) before extracting them for structure determination. Reference-free 2D classification was performed in ROME (Wu et al., 2016), whereas unsupervised 3D classification and high-resolution refinement was performed in RELION 1.3 (Scheres, 2012a, b). Reported resolutions are based on the gold-standard FSC 0.143 criterion, and FSC curves were corrected for the effects of a soft mask. Prior to visualization, all density maps were corrected for the modulation transfer function (MTF) of the detector, and then sharpened by applying a negative B-factor that was estimated using automated procedures (Tang et al., 2007). Local resolution variations were estimated using ResMap (Kucukelbir et al., 2014). Details in data analysis are provided in Supplemental Experimental Procedure.

### Atomic Model Building and Refinement

Direct rigid-body fitting of crystal structures of yeast Rpn2, Rpn6, Rpn9, Rpn10 and Rpn12 monomers and Rpn11-Rpn8 dimer or into our cryo-EM map suggests an excellent agreement in the secondary structural elements between our cryo-EM structure and the crystal structures. The initial atomic modelling of non-AAA subcomplex was based on the atomic structure of human 26S proteasome (Chen et al., 2016). Refinement was first carried out on Rpn subunits and 6 Rpt subunits with CP in real space using Phneix with secondary structure and geometry restraints to prevent over-fitting. Each part of cryo-EM map was segmented by Chimera. The cryo-EM map and the atomic model were placed into a pseudo-unit cell and the refinement was performed in Phenix (Adams et al., 2010) in Fourier space using both amplitudes and phases, with restraints by non-crystallographic symmetry. The final refinement statistics are shown in Table S1. Pseudo-atomic models of T_1_ −T_3_, except the AAA domain, Rpn1 and gankyrin, were based on the atomic model of non-AAA subcomplex. Pseudo-atomic models of Rpn1-p28-AAA subcomplex (T_A1_-T_A7_) were based on atomic structure of human 26S proteasome (Chen et al., 2016) and crystal structure of gankyrin (Nakamura et al., 2007a) (PDB accession code: 2DWZ).

### Structural Analysis and Visualization

Solvation energy and interaction surface for each model were calculated using PISA (Krissinel and Henrick, 2007). Total interaction surface or free energy was calculated by summing up the contributions from each interaction pairs. Bias due to truncation of the termini of some subunits was compensated after the calculation. Subunit-subunit interaction networks or transition networks were visualized using Pajek (http://mrvar.fdv.uni-lj.si/pajek/). Numeric processing was carried out in Matlab. The scripts are available upon request.

## SUPPLEMENTAL INFORMATION

Supplemental Information includes Supplemental Experimental Procedures, eight figures, three tables, and can be found with this article online.

## AUTHOR CONTRIBUTIONS

Y.L., S.C. and S.S. purified proteasome and performed biochemical experiments. Y.L. prepared the specimen for data collection and developed the unfolding assay. Y.M. collected cryo-electron microscopy data. S.C. performed initial reconstruction. Y.L., J.W. and Y.M. designed computational strategy for data processing. J.W., Y.L. and Y.B.M. processed the data and refined the cryo-EM maps. Y.L. and Y.M. built the initial atomic model. S.C. and Y.M. refined the atomic model for the near-atomic resolution structure and built the pseudo-atomic models for the medium-resolution structures. All authors contributed to data analysis and manuscript preparation.

## ACKNOWLEDGEMENTS

The authors thank J. Jackson and T. Song for assistance in maintaining the high-performance computing system; D. Yu for assistance in drift correction; Dr. Horwich for a gift of the ClpA and GroEL constructs. This work was funded in part by a grant from NIGMS, GM02675 (M.K.), by a grant of the Thousand Talents Plan of China (Y.M.), by an Intel academic grant (Y.M.), by the research funds at Peking-Tsinghua Center for Life Science at Peking University (Q. O.), by a grant from National Natural Science Foundation of China 91530321 (Y.M., Q.O.), and by the NIH grant R01 GM095526 (D.F.). The cryo-EM experiments were performed in part at the Center for Nanoscale Systems at Harvard University, a member of the National Nanotechnology Coordinated Infrastructure Network (NNCI), which is supported by the National Science Foundation under NSF award no. 1541959. The cryo-EM facility was funded through the NIH grant AI100645, Center for HIV/AIDS Vaccine Immunology and Immunogen Design (CHAVIID). The data processing was performed in part in the Sullivan supercomputer, which is funded in part by a gift from Mr. and Mrs. Daniel J. Sullivan, Jr.

